# Enhanced efficacy of a specific HDAC3 inhibitor in combination with 5-Azacitidine against diffuse large B-cell lymphoma

**DOI:** 10.1101/2025.04.21.648509

**Authors:** Calixte Calaud, Emilio Serrano-Lopez, Leonie A. Cluse, Izabela Todorovski, Thomas Mardivirin, Matt Teater, Abdelillah Maarad, Noura Tawfic, Sophia Rutaquio, Peter J. Fraser, Killian Guiraud, David Yoannidis, Emeline Kerreneur, Arnaud Jacquel, Ari M. Melnick, Ricky W. Johnstone, Pilar M. Dominguez

## Abstract

Diffuse large B-cell lymphoma (DLBCL) refers to an aggressive lymphoma that arises from germinal center (GC) B-cells, which differentiate into plasma cells (PC) to produce high affinity antibodies. 40% of DLBCL patients relapse or are refractory to the conventional immunochemotherapy treatment, usually with fatal consequences. Therefore, there is an unmet critical need to find more targeted therapies for DLBCL. DLBCL are characterized by profound alterations in the epigenome, which are correlated with poor survival. While epigenetic therapies are used as anti-cancer treatments, their full potential has not been achieved, mainly because of their limited efficacy when used as monotherapies and recurrent side effects associated with their low specificity. The abnormal epigenetic landscape of DLBCL tumors is associated with a blockade in GC exit and differentiation programs, which are regulated by the transcription factor BCL6. This aberrant repression of BCL6-target genes is mediated at least by two epigenetic mechanisms: 1) increased DNA methylation and 2) loss of acetylation of the lysine 27 of histone 3 (H3K27ac)-through recruitment of histone deacetylase 3 (HDAC3). Therefore, we investigated the efficacy against DLBCL of a novel combinatorial epigenetic therapy using the hypomethylating agent (HMA) 5-Azacitidine (5-Aza) and a specific HDAC3 inhibitor (HDAC3i). We found that treatment with 5-Aza and HDAC3i had a potent synergistic anti-tumor activity *in vitro* and *in vivo*, which was superior to the effect of each single drug or 5-Aza combined with non-specific HDACi and, importantly, was not associated with toxicity in normal cells. We also demonstrated that the combined 5-Aza and HDAC3i treatment induced the epigenetic remodeling of DLBCL cells, which resulted in a more potent re-expression of PC differentiation genes, including *XBP1* and *ATF4*, compared to each drug used as single agents. Our results highlight the importance of targeting multiple layers of the epigenome to maximize the efficacy of epigenetic-based therapies.

## INTRODUCTION

Diffuse large B-cell lymphoma (DLBCL), a blood cancer that originates from germinal center (GC) B cells, is the most common type of lymphoma in adults. DLBCL are categorized as GC B cell-like (GCB), activated B cell-like (ABC) or unclassified based on their cell of origin (1,2). Due to its aggressiveness, only 60% DLBCL patients respond to the first-line immuno-chemotherapy consisting of the monoclonal antibody rituximab and four drugs (cyclophosphamide, doxorubicin, vincristine, and prednisone), known as R-CHOP (3). The other 40% of DLBCL patients present relapsed or refractory (R/R) disease, which is associated with poor prognosis and a median overall survival of only 6,3 months (4). Therefore, there is a clinical need to find improved therapies to treat DLBCL.

DLBCL is characterized by alterations in the epigenome, including recurrent somatic mutations in genes encoding epigenetic enzymes and chromatin modifiers as well as aberrant DNA methylation patterns compared to normal GC B-cells (5). Importantly, multiple studies from us and others have shown that epigenetic dysregulation contributes to DLBCL initiation and progression (6). This knowledge has led to the development during the last two decades of epigenetic-based therapies to treat DLBCL, including inhibitors of DNA methyltransferases (DNMT) and histone deacetylases (HDAC) (7).

The DNMT inhibitors (DNMTi) 5-Azacitidine (5-Aza) and Decitabine are approved by the US Food and Drug Administration (FDA) and the European Medicines Agency, and they are currently used to treat myeloid diseases. In DLBCL, preclinical studies have showed that lymphoma cells exposed to low doses of 5-Aza for a prolonged time become sensitive to chemotherapy through the hypomethylation and subsequent reactivation of genes as *SMAD1*, which is hypermethylated in chemoresistant DLBCL cells (8). An associated phase I clinical trial consisting of 5-Aza prior to R-CHOP resulted in complete responses (CR) in 10/11 high-risk DLBCL patients (8). A more recent phase I clinical trial of oral 5-Aza (CC-486) plus R-CHOP in 59 intermediate-to high-risk DLBCL patients confirmed the efficacy of 5-Aza priming before chemotherapy with an overall response rate (ORR) of 94,9% and CR achieved in 88% patients (9).

Despite these promising results, 5-Aza treatment at the effective doses was associated with hematological toxicities such as neutropenia (9), as is the case with pan-HDAC inhibitors (HDACi), which target multiple isoforms. Previous studies evaluating the efficacy of pan-HDACi in monotherapy to treat DLBCL have shown limited clinical responses and frequent off-target effects (10). A potential strategy to overcome the limitations of epigenetic therapies when used as monotherapies, increasing their efficacy while reducing their undesired toxic side effects, is through the design of new combinatorial therapies. The combination of Decitabine plus HDACi showed a synergistic effect in both GCB- and ABC-DLBCL cell lines, primary DLBCL cells *in vitro* and murine xenograft models (11). A more recent study combining 5-Aza and the HDACi Vorinostat confirmed their synergistic activity against DLBCL, although the accompanying phase Ib trial with 18 DLBCL patients revealed low clinical benefit and associated toxicities in patients treated with the combinatorial therapy (12).

Next-generation selective HDACi are currently in development and show promise to improve the safety profiles of this type of epigenetic inhibitors (13). In this regard, we previously showed that during the GC reaction HDAC3 is recruited by BCL6 in GC B-cells to induce the repression of BCL6-target genes, which are re-expressed upon GC exit by the histone acetyl transferase CREBBP to allow the normal differentiation of GC B-cells into plasma cells (PC) o memory B-cells (14-16). Moreover, we demonstrated that the selective inhibition of HDAC3 was effective against DLBCL, especially in tumors with mutations in the histone acetyltransferases (HAT) *CREBBP* (17) or TET2-deficient lymphomas (18).

Due to this essential role of HDAC3i in GC biology and lymphomagenesis, we hypothesized that the anti-tumor effects of hypomethylating agents (HMA) can be enhanced by the inhibition of HDAC3. Here we investigate the therapeutic efficacy of 5-Aza in combination with a specific HDAC3i against DLBCL, compared to each single agent or 5-Aza plus pan-HDACi.

## RESULTS

### Specific HDAC3 inhibition synergizes with 5-Aza against DLBCL cells

To analyze the effect on DLBCL cells of inhibiting DNMT in combination with inhibition of HDAC3, we used 5-Aza and a specific HDAC3i (KDAc-001) (19). We treated DLBCL cells with 5-Aza daily for 5 days (day 0–day 4) and at days 2 and 4 we added the HDAC3i (**Fig. 1A**). This sequential treatment allows 5-Aza to be incorporated into the DNA and exert its function before cells undergo growth arrest due to HDAC3i treatment. In addition, we used low doses of 5-Aza (50-200nM) to induce demethylation of DNA without causing cytotoxicity. In MD901 we observed no induction of cell death based on propidium iodide (PI) staining after single treatment with 5 or 10μM HDAC3i and a moderate increase in the percentage of PI^+^ cells after treatment with 200nM 5-Aza alone (**Fig. 1B**). Notably, when cells were treated with both drugs we observed a cytotoxic dose-dependent effect at day 7 (**Fig. 1B**). We found similar results when OCI-Ly7 cells were treated with 5-Aza and HDAC3i (**Fig. 1B**). We analyzed the mechanism of cell death, and we found a significant increase in late apoptotic cells (Annexin V^+^ PI^+^) in the 5-Aza+HDAC3i condition compared to MD901 and OCI-Ly7 cells treated with vehicle, 5-Aza or HDAC3i (**Fig. 1C** and **Supplementary Fig. S1A**). In line with these results, the level of cleaved Caspase-3 was significantly higher in combo-treated MD901 and OCI-Ly7 cells than in cells treated with 5-Aza or HDAC3i alone (**Fig. 1D** and **Supplementary Fig. S1B**). Then we investigated if the treatment had a cytostatic effect. First, we analyzed the total number of cells; we found a decrease in the cell number after treatment with 10μM HDAC3i and a more clear reduction when cells were treated with 5-Aza alone, in line with the anti-proliferative activity described for other HMA on DLBCL (12,20) (**Fig. 1E**). Significantly, the combination of 5-Aza+HDAC3i resulted in a greater dose-dependent reduction in the total cell number compared to cells treated with each single agent (**Fig. 1E** and **Supplementary Fig. S1C**). To further investigate the anti-proliferative effect of the combinatorial treatment we labeled the cells with CellTrace Violet (CTV) and analyzed the number of divisions. Treatment with 5-Aza alone or in combination with HDAC3i delayed proliferation at day 2 (**Fig. 1F** and **Supplementary Fig. S1D**). At day 4 we observed that the blockade in proliferation was greater in the 5-Aza+HDAC3i condition than in cells only treated with 5-Aza or HDAC3i (**Fig. 1F and 1G, and Supplementary Fig. S1C**). Importantly, we confirmed that the anti-lymphoma effect of combining 5-Aza+HDAC3i was synergistic in both MD901 and OCI-Ly7 cells (**Fig. 1H**).

**FIGURE 1:**
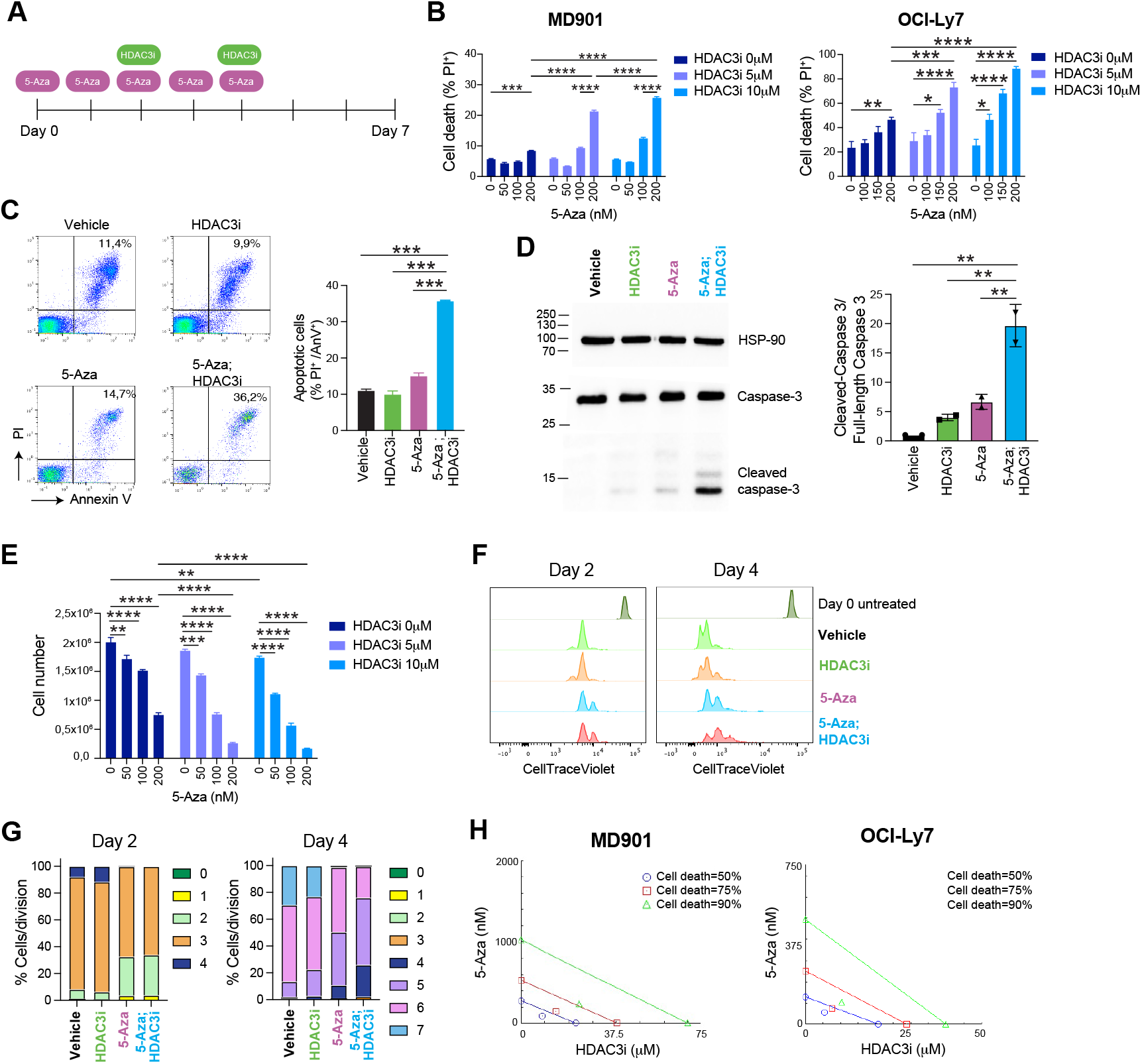
5-Aza and HDAC3i synergistic effect results in increased cell death and reduced proliferation in DLBCL cells. (**A**) Schematic representation of the treatment of DLBCL cells with 5-Aza and HDAC3i. (**B**) Percentage of PI^+^ cells at day 7 after treatment of MD901 and OCI-Ly7 cells with vehicle, HDAC3i, 5-Aza or 5-Aza+HDAC3i. (**C**) Representative density plot and quantification of Annexin V^+^ PI^+^ cells at day 7 after treatment of MD901 cells with vehicle, HDAC3i, 5-Aza or 5-Aza+HDAC3i. (**D**) Representative western blot of full-length Caspase 3 and cleaved-Caspase 3, and quantification relative to control at day 5 after treatment of MD901 cells with vehicle, HDAC3i, 5-Aza or 5-Aza+HDAC3i. (**E**) Total cell number of MD901 cells at day 7 after treatment with vehicle, HDAC3i, 5-Aza or 5-Aza+HDAC3i. (**F, G**) CTV assay at day 2 and day 4 after treatment of MD901 cells with vehicle, HDAC3i, 5-Aza or 5-Aza+HDAC3i; (**F**) representative density plot and (**G**) percentage of cells per division. (**H**) Isobologram analysis of MD901 and OCI-Ly7 cells at day 7 after treatment with 5-Aza and HDAC3i.

### Treatment of DLBCL cells with 5-Aza in combination with HDAC3i induces specific transcriptional signatures associated with differentiation into plasma cells

To identify the molecular mechanisms responsible for the synergistic effect of 5-Aza and HDAC3i on DLBCL, we analyzed the impact of the combinatorial epigenetic treatment on the transcriptional profile of lymphoma cells by RNA-sequencing (RNA-seq) at day 5. Principal component analysis showed a clear difference in transcriptional profiles between DLBCL cells treated with vehicle, HDAC3i, 5-Aza and 5-Aza+HDAC3i (**Supplementary Fig. S2A**). Next, we performed a supervised analysis and identified 1417 DEG between 5-Aza+HDAC3i and vehicle, 879 DEG between 5-Aza and vehicle, and 181 differentially expressed genes (DEG) between HDAC3i and vehicle (log fold change (logFC)>1, P<0.05; **Fig. 2A and 2B** and **Supplementary Table S1**). Pathway analysis was performed using gene set enrichment analysis (GSEA) using hallmark gene sets. We observed a significant enrichment for upregulation of genes involved in the unfolded protein response (UPR) and the epithelial-mesenchymal transition in cells treated with the 5-Aza+HDAC3i combination (**Fig. 2C**, left and **Supplementary Table S2**). Treatment with 5-Aza alone resulted in significant enrichment for upregulation of only 4 pathways, which were related to protein secretion, TGF beta (TGFβ) and IL-6 signaling (**Fig. 2C**, middle and **Supplementary Table S2**), whereas treatment with HDAC3i induced significant positive enrichment of immune and inflammatory pathways, in line with previous studies (17) (**Fig. 2C**, right and **Supplementary Table S2**). We found fewer number of significantly enriched gene sets among downregulated genes, suggesting that the epigenetic drugs preferentially induce transcriptional activation; the pathways with negative enrichment scores included MYC targets, oxidative phosphorylation and DNA repair for cells treated with HDAC3i or 5-Aza+HDAC3i, and E2F targets for 5-Aza-treated cells (**Supplementary Fig. S2B** and **Supplementary Table S3**). We further characterized the effect of the combinatorial therapy at the transcriptional level using a comprehensive database of gene sets relevant for GC B cells and DLBCL, derived from our previous work and others. This analysis revealed significant positive enrichment for BCL6 target genes and pathways involved in PC differentiation, including upregulation of the gene encoding for the key transcription factor (TF) XBP1, after treatment with 5-Aza+HDAC3i (**Fig. 2D** and **Supplementary Table S4**). This aligns with the significant UPR enrichment observed using Hallmark datasets (**Fig. 2C**), since activation of the UPR is a critical process of PC function and is mediated by XBP1 (21). Conversely, the analysis of HDAC3i-treated cells showed significant enrichment for upregulation of genes induced by CD40 signaling, including major histocompatibility (MHC) genes, as previously shown (17) (**Fig. 2D** and **Supplementary Table S4**), whereas 5-Aza induced significant enrichment for upregulation of BCL6 target genes and genes repressed in DLBCL cells resistant to chemotherapy (8) (**Fig. 2D** and **Supplementary Table S4**). We confirmed by RT-PCR that the treatment with 5-Aza+HDAC3i increased the expression of genes upregulated in PC: *XBP1, TNFRSF17, ATF4, FNDC3B, CD38* and *EDEM1* (22,23) (**Fig. 2E**). In addition, we used the CollecTRI meta-resource to infer changes in TF activity based on the differentially expressed transcriptomes (24) and we found that ATF4 had the highest increased activity in DLBCL cells treated with 5-Aza+HDAC3i (**Fig. 2F** and **Supplementary Table S4**), confirming the results of the GSEA showing activation of the UPR. CTCFL and FOXO3 were found to have increased activity in cells treated with 5-Aza; and cells treated with HDAC3i showed increased activity in IRF1 and CIITA, which are IFN-related TF (**Fig. 2F** and **Supplementary Table S4**), in line with previous work (17). These results suggest that the synergistic effect of 5-Aza and HDAC3i against DLBCL when used in combination (**Fig. 1E**) is linked to the induction of terminal differentiation towards PC.

**FIGURE 2:**
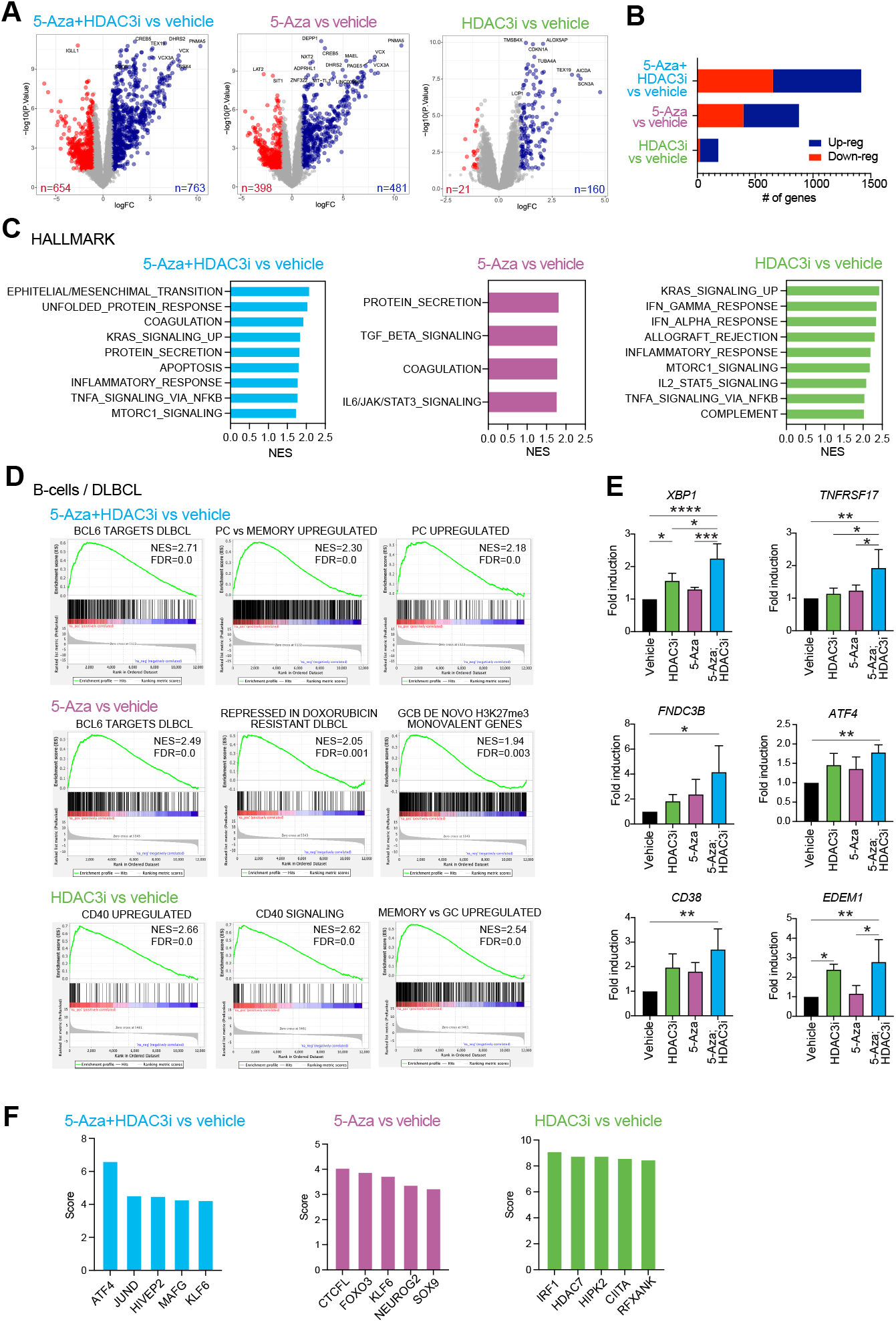
5-Aza and HDAC3i induce different transcriptional signatures in DLBCL cells as single agents than in combination. (**A**) Volcano plot of the DEG (logFC > 1; *P*val < 0.05) between drug-treated (5-Aza+HDAC3i, 5-Aza, HDAC3i) and vehicle-treated DLBCL cells at day 5. (**B**) Quantification of the DEG shown in **A**. (**C**) Gene set enrichment analysis using the Hallmark gene sets. The top 10 more significant gene sets in each treatment are shown. (**D**) Gene set enrichment analysis using B-cell/DLBCL-associated gene sets. (**E**) Expression by qRT-PCR of plasma cell-specific genes on treated DLBCL cells collected at day 5 (n=2; \**P* < 0.05; \*\**P* < 0.01, \*\*\**P* < 0.001, \*\*\*\**P* < 0.0001). (**F**) Top 5 TF inferred to be activated in each condition.

### The 5-Aza+HDAC3i combination induces epigenetic changes in DLBCL

We analyzed the genome-wide epigenetic changes induced by the treatment, focusing on the epigenetic marks targeted by each drug: DNA methylation (5-Aza) and histone acetylation (HDAC3i). The analysis of the methylome of DLBCL cells, performed through reduced representation bisulfite sequencing (RRBS), showed a well-defined separation between conditions with or without 5-Aza (**Supplementary Fig. S3A**). The analysis of differentially methylated cytosines (DMC) revealed massive hypomethylation in cells treated with 5-Aza or 5-Aza+HDAC3i compared to vehicle-treated cells (methylation difference>25%; FDR<0.05; **Fig. 3A** and **Supplementary Fig. S3B**). Treatment with HDAC3i induced few changes in the methylome compared to vehicle (**Fig. 3A** and **Supplementary Fig. S3B**). We overlapped the hypomethylated DMC (hypo-DMC) between 5-Aza vs vehicle and 5-Aza+HDAC3i vs vehicle and we found that the 97% were shared (**Fig. 3B**), distributed mostly in introns and intergenic regions (**Supplementary Fig. S3C**).

**FIGURE 3:**
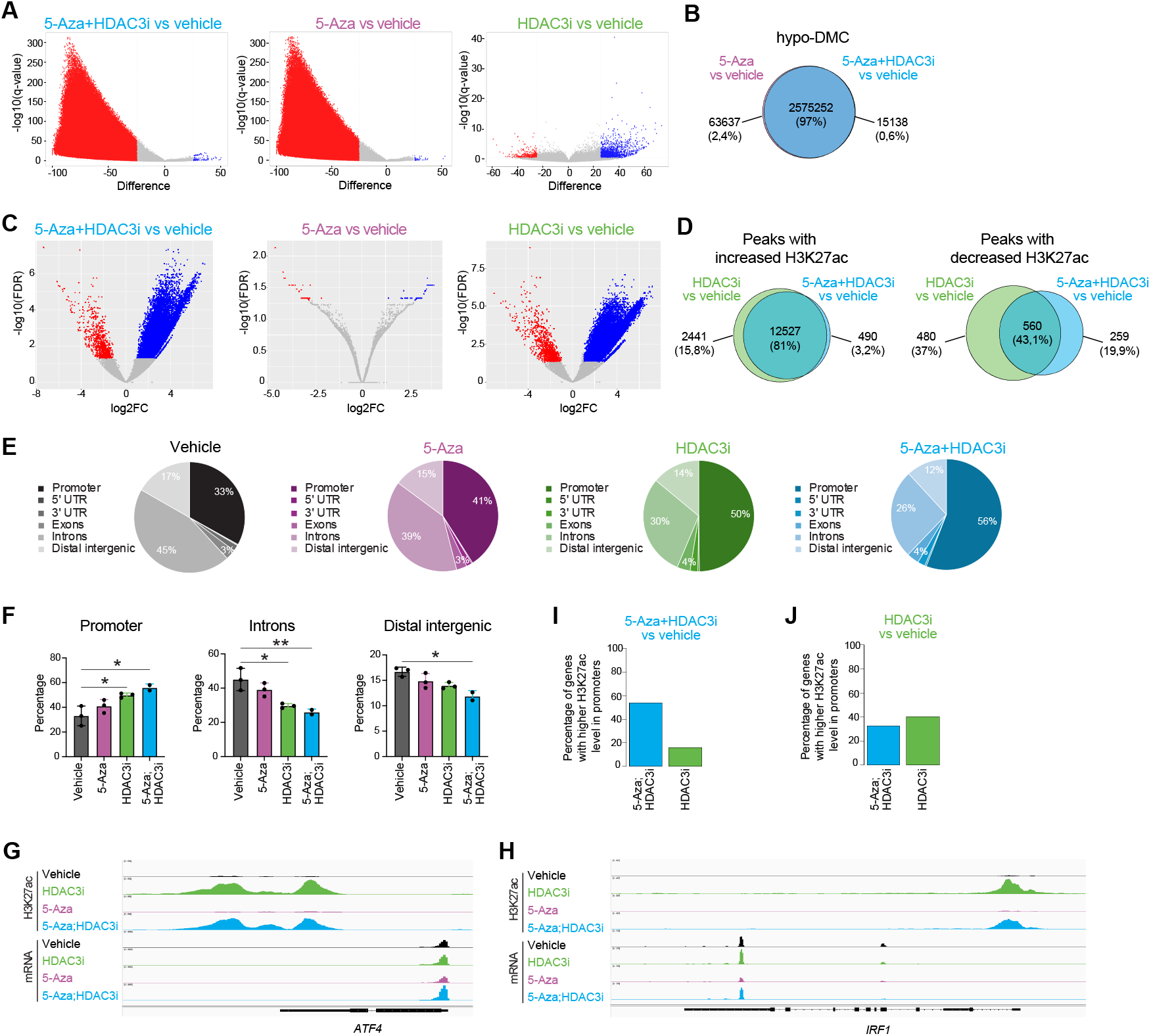
Epigenetic changes induced in DLBCL cells after treatment with 5-Aza and/or HDAC3i. (**A**) Volcano plot of the DMC (25% methylation difference; FDR < 0.05) between drug-treated (5-Aza+HDAC3i, 5-Aza, HDAC3i) and vehicle-treated DLBCL cells at day 5. (**B**) Overlap of the hypo-DMC between 5-Aza+HDAC3i vs vehicle and 5-Aza vs vehicle. Volcano plot of the differential H3K27ac peaks (logFC > 1; FDR < 0.05) between drug-treated (5-Aza+HDAC3i, 5-Aza, HDAC3i) and vehicle-treated DLBCL cells at day 5. (**D**) Overlap of the increased and decreased H3K27ac peaks between 5-Aza+HDAC3i vs vehicle and HDAC3i vs vehicle. (**E**) Genomic distribution of the H3K27ac peaks in each condition. (**F**) Comparison of the genomic distribution of the H3K27ac peaks in each condition. (**G**) Read-density tracks of normalized H3K27ac ChIP-seq and RNA-seq at the *ATF4* locus. (**H**) Read-density tracks of normalized H3K27ac ChIP-seq and RNA-seq at the *IRF1* locus. (**I**) Percentage of genes positively enriched in 5-Aza+HDAC3i vs vehicle with >25% higher promoter H3K27ac intensity in 5-Aza+HDAC3i compared to HDAC3i (blue) or viceversa (green). (**J**) Percentage of genes positively enriched in HDAC3i vs vehicle with >25% higher promoter H3K27ac intensity in 5-Aza+HDAC3i compared to HDAC3i (blue) or viceversa (green).

Chromatin immunoprecipitation followed by sequencing (ChIP-seq) was used to determine the levels of H3K27 acetylation (H3K27ac). Principal component analysis showed a clear separation between the conditions with or without HDAC3i (**Supplementary Fig. S3D**). The subsequent analysis of differential peaks compared to vehicle identified higher number of peaks with increased acetylation than reduced acetylation in the conditions treated with HDAC3i or 5-Aza+HDAC3i (logFC>1; FDR<0.05; **Fig. 3C** and **Supplementary Fig. S3C**). Treatment with 5-Aza induced changes in H3K27ac but most of them were under the significance threshold (logFC>1; FDR<0.05; **Fig. 3C** and **Supplementary Fig. S3C**). Further analysis of the differential peaks gaining H3K27ac revealed that 81% were shared when comparing HDAC3i vs vehicle and 5-Aza+HDAC3i vs vehicle (**Fig. 3D**), which could be a reflection of the direct effect of HDAC3 inhibition. In contrast, only 43% of the peaks losing H3K27ac were shared between HDAC3i vs vehicle and 5-Aza+HDAC3i vs vehicle, a result that could be explained by the indirect, and therefore more random, effects of HDAC3i (**Fig. 3D**).

Next, we annotated the peaks into genomic features. We observed a gradual increase in the percentage of H3K27ac peaks in promoters: vehicle=33%, 5-Aza=41%, HDAC3i=50% and 5-Aza+HDAC3i=56%, which correlated with a decrease in the percentage of peaks both in introns: vehicle=45%, 5-Aza=39%, HDAC3i=30% and 5-Aza+HDAC3i=26% and distal intergenic regions: vehicle=17%, 5-Aza=15%, HDAC3i=14% and 5-Aza+HDAC3i=12% (**Fig. 3E**). The compilation of this data showed significantly higher acetylation at promoters and reduced acetylation at introns after HDAC3i or 5-Aza+HDAC3i treatment compared to vehicle, and significantly reduced acetylation at distal intergenic regions in the 5-Aza+HDAC3i vs vehicle condition (**Fig. 3F**). To investigate if these differences in H3K27ac at promoters could contribute to the distinct transcriptional profiles between HDAC3i-treated and combo-treated cells, we overlapped the H3K27ac and RNA-seq profiles. First, we focused on the key TF identified on each condition: *ATF4* (5-Aza+HDAC3i) and *IRF1* (HDAC3i). We observed an increase in the level of H3K27ac at the promoter of *ATF4* after 5-Aza+HDAC3i treatment compared to HDAC3i alone, which correlated with increased transcription in the combo-treated cells (**Fig. 3G**). Conversely, the promoter of *IRF1* showed a higher level of H3K27ac after treatment with HDAC3i than with 5-Aza+HDAC3i, and this correlated with higher upregulation of *IRF1* in the HDAC3i-treated cells compared to combo-treated cells (**Fig. 3H**). We expanded this analysis to all the genes that were positively enriched in each condition based on the GSEA (**Fig. 2C**). We found that the majority (54%) of genes that are uniquely transcriptionally activated after 5-Aza+HDAC3i have at least 25% higher H3K27ac at promoters in 5-Aza+HDAC3i than in HDAC3i alone, including *ATF4, EDEM1* and *CD38* (**Fig. 3I**). When analyzing the genes positively enriched after HDAC3i treatment, the differences were smaller, i.e. only 40% had higher promoter intensity in HDAC3i and 33% showed higher H3K27ac at promoters after treatment with 5-Aza+HDAC3i (**Fig. 3J**). Overall, these results suggest that 5-Aza priming enhances the transcriptional differentiation program that is initiated by the inhibition of HDAC3, through the redistribution and increased intensity of H3K27ac specially at BCL6-target genes.

### Superior therapeutic efficacy of 5-Aza+HDAC3i against DLBCL compared to 5-Aza+pan-HDACi

Next, we tested if the anti-lymphoma activity of 5-Aza+HDAC3i was superior to 5-Aza combined with pan-HDACi: Panobinostat or Vorinostat. First, we performed dose titrations of each HDACi in DLBCL cells to select equivalent non-toxic concentrations of these three inhibitors when used as single agents: 2nM Panobinostat, 100nM Vorinostat and 10μM HDAC3i (**Supplementary Fig. S3A**). Then, we combined each HDACi with 5-Aza. Treatment of DLBCL cells with the combination consisting of 5-Aza+HDAC3i resulted in greater induction of cell death (42% PI^+^ cells) compared to 5-Aza+Panobinostat (24% PI^+^ cells) and 5-Aza+Vorinostat (23% PI^+^ cells) (**Fig. 4A**). We also found differences in proliferation, with a significant lower cell number after treatment with 5-Aza+HDAC3i compared to cells treated with 5-Aza alone, 5-Aza+Panobinostat or 5-Aza+Vorinostat (**Fig. 4B**).

**FIGURE 4:**
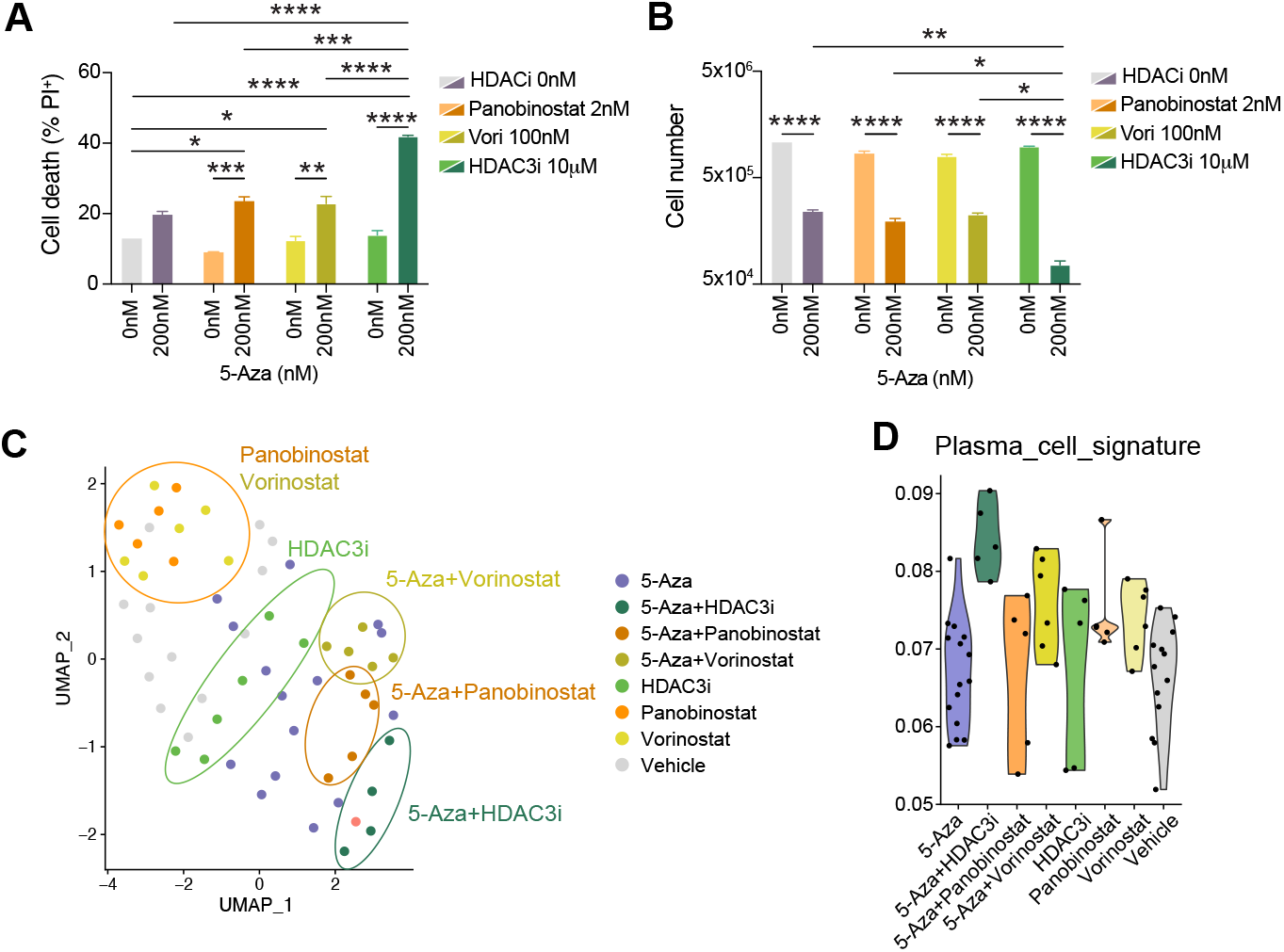
Superior efficacy of 5-Aza+HDAC3i against DLBCL compared to 5-Aza+pan-HDACi. (**A**) Percentage of PI^+^ cells at day 7 after treatment of MD901 cells with vehicle or 5-Aza in combination with HDAC3i, Panobinostat or Vorinostat. (**B**) Total cell number of MD901 cells at day 7 after treatment with vehicle or 5-Aza in combination with HDAC3i, Panobinostat or Vorinostat. (**C**) UMAP plot for the transcriptome data obtained at day 4 after treatment of MD901 cells with vehicle or 5-Aza in combination with HDAC3i, Panobinostat or Vorinostat. Enrichment score for the plasma cell signature in each condition.

To simultaneously characterize the phenotypic and transcriptional effects of the different treatments we used Multiplexed Analysis of Cells sequencing (MAC-seq) (25,26), which is based on the Digital RNA with perturbation of Genes (DRUG-seq) method (27). This approach allowed us to perform in one single assay a direct comparison of the therapeutic efficacy and transcriptional changes induced by the three combinatorial therapies. We confirmed in this miniaturized high-throughput setting that the treatment with 5-Aza+HDAC3i had a stronger therapeutic effect against DLBCL, with higher percentage of PI^+^ cells than in the conditions treated with 5-Aza+Panobinostat or 5-Aza+Vorinostat (**Supplementary Fig. S3B**). Using the same assay, we investigated the molecular mechanisms driving the stronger cytotoxic and cytostatic effects induced by the 5-Aza+HDAC3i combination. Uniform Manifold Approximation and Projection for Dimension Reduction (UMAP) analysis showed that the conditions consisting of single treatment with Panobinostat and Vorinostat clustered closely together, while cells treated with HDAC3i alone resulted in a different transcriptomic profile, situated between single pan-HDACi and combo conditions (**Fig. 4C**). Moreover, the combinations with 5-Aza clustered separately than their single counterparts, with the 5-Aza+HDAC3i combination showing a more different transcriptomic response compared to 5-Aza+Panobinostat or 5-Aza+Vorinostat (**Fig. 4C**). Then, we overlapped the PC signature identified in cells treated with 5-Aza plus HDAC3i (**Fig. 2**). This analysis showed that the combination of 5-Aza+HDAC3i resulted in higher upregulation of genes linked to PC differentiation compared to the other combinations or single treatments (**Fig. 4C**).

### Translational potential of the 5-Aza+HDAC3i combination as a new therapy for DLBCL

After confirming the anti-lymphoma activity of the new combinatorial epigenetic treatment consisting of 5-Aza and HDAC3i, we aimed to investigate its safety in non-tumor cells. We treated peripheral blood mononuclear cells (PBMC) with 5-Aza+HDAC3i, 5-Aza, HDAC3i or vehicle. We observed no toxic effects of the combinatorial therapy on T cells (**Fig. 5A**) and only a slight toxicity on B cells at the highest concentration of both drugs (**Fig. 5B**), which was lower compared to the effect of the less specific HDACi Panobinostat and comparable to the effect of Vorinostat (**Fig. 5A** and **5B**).

**FIGURE 5:**
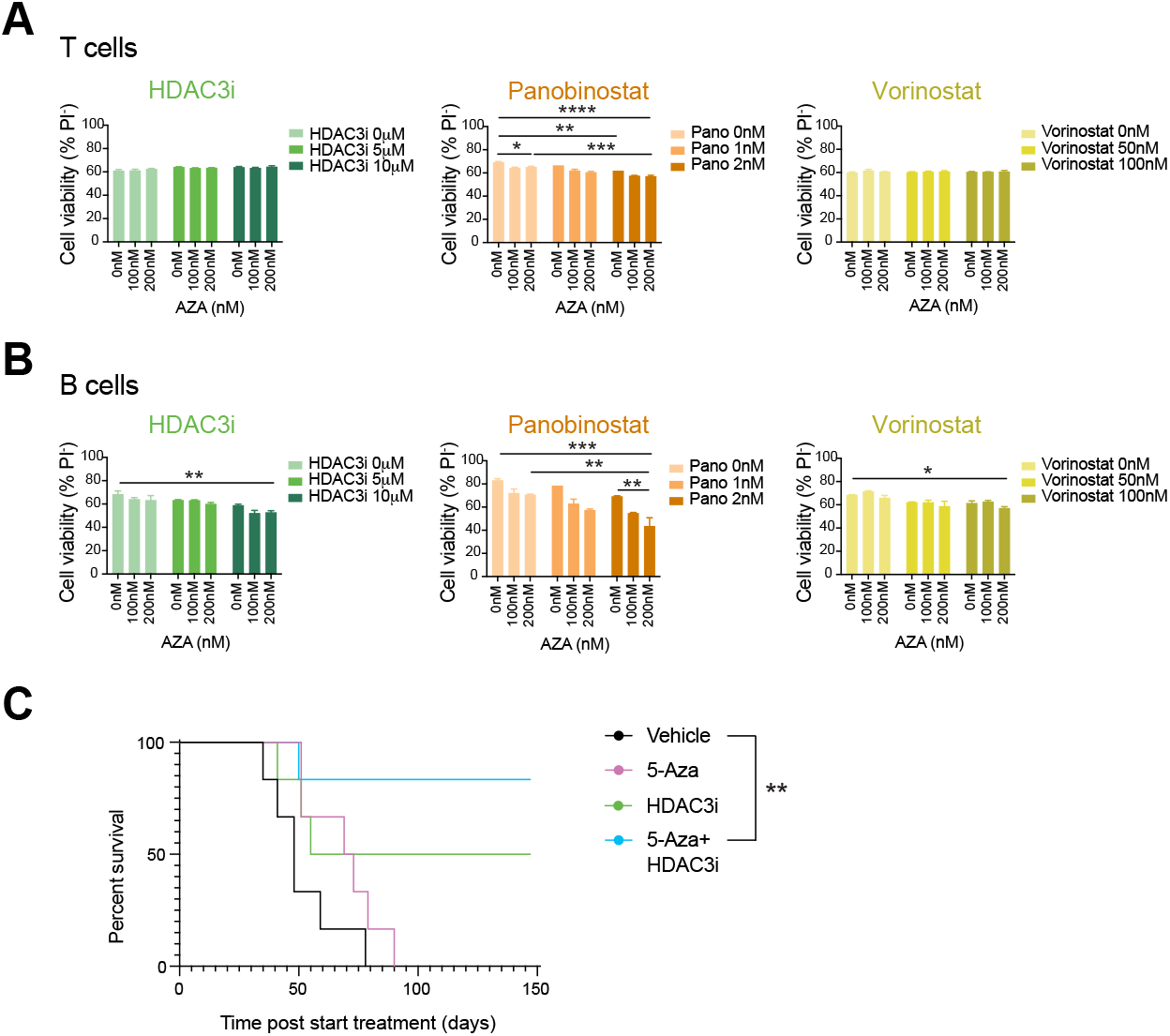
Translational potential of the 5-Aza+HDAC3i combination as a new therapy for DLBCL. (**A**) Percentage of PI^+^ cells at day 7 after treatment of T cells with vehicle or 5-Aza in combination with HDAC3i, Panobinostat or Vorinostat. (**B**) Percentage of PI^+^ cells at day 7 after treatment of B cells with vehicle or 5-Aza in combination with HDAC3i, Panobinostat or Vorinostat. (**C**) Kaplan–Meier survival curves of mice transplanted with OCI-Ly7 cells and treated with either vehicle, 5-Aza, HDAC3i or 5-Aza+HDAC3i (n=6 mice/group).

Finally, we investigated the *in vivo* efficacy of the combination compared to treatment with each single drug. We generated xenografts transplanting OCI-Ly7 into immunodeficient NSG mice, intraperitoneally to obtain systemic disease. Treatment with 5-Aza+HDAC3i induced a significant survival advantage compared to vehicle (**Fig. 5C**). Moreover, 80% mice treated with the combination survived for 150 days compared to only 50% mice surviving after treatment with HDAC3i alone and 0% in the case of mice receiving 5-Aza (**Fig. 5C**). In addition, we did not observe significant changes in body weight in the combo-treated mice throughout the duration of the treatment, indicating that the combination has an *in vivo* safe profile (**Supplementary Fig. S4**).

## DISCUSSION

Since the pathogenesis of DLBCL is strongly linked to epigenetic abnormalities, including DNA methylation patterning and histone modifications, different classes of epigenetic therapies have been developed to treat this type of lymphoma (10). However, although promising, these therapeutic approaches are not broadly implemented in the clinic for DLBCL patients, mainly due to their limited efficacy when used as single agents. Here we show that the treatment with 5-Aza in combination with HDAC3i results in synergistic enhanced cytotoxicity and lower proliferative capacity in DLBCL cells. Our findings are consistent with previous studies investigating the anti-tumor effect of combining HMA and HDACi in DLBCL (11,12), but 5-Aza plus a specific HDAC3i have a more potent anti-lymphoma activity compared to pan-HDACi without affecting the viability of healthy lymphocytes. Importantly, we confirmed the superior efficacy compared to single agents of this precise combination of epigenetic inhibitors using a DLBCL xenograft model of systemic disease, highlighting the improved therapeutic potential of our proposed epigenetic combinatorial therapy.

Mechanistically, we found that 5-Aza and HDAC3i in combination induced a different transcriptomic profile than as monotherapies, supporting the existence of a cooperative mechanism of action leading to the observed synergistic cytotoxic effect. In this regard, the combinatorial therapy resulted in activation of genes involved in the UPR and PC differentiation, including *XBP1* and *ATF4*. The differentiation program of GC B-cells into PC requires the production of immunoglobulin chains through the endoplasmic reticulum (ER), which leads to activation of the UPR (28); however, prolonged UPR results in the elimination of the stressed cells by apoptosis (29). Therefore, a possible explanation for our results is that the activation of the PC differentiation pathway mediated by the combinatorial therapy leads to terminal ER stress and, since the cells are unable to restore homeostasis, the maintained activation of the UPR ultimately induces apoptosis (28).

Since impaired terminal differentiation and inability to exit the GC reaction are features of the transformation of GC B-cells into DLBCL, therapies promoting cell differentiation are expected to disrupt the malignant properties of lymphoma cells, as is the case for myeloid malignancies (30). Interestingly, epigenetic-targeted drugs including HMA and HDACi are among the most effective therapies to relieve the differentiation blockade of leukemia cells (30). In this regard, we previously demonstrated that treatment with the HDACi Panobinostat induced differentiation of acute myeloid leukemia cells (31,32). In the present study, the PC transcriptional signature observed when combining 5-Aza and HDAC3i aligns with the findings of two recent studies showing a synergistic effect of targeting DOT1L and EZH2 against GCB-DLBCL cells, which is associated with the differentiation towards a PC-like state (33,34). Combined inhibition of EZH2 and DOT1L resulted in upregulation of BCL6-target genes associated with IFN-mediated responses, antigen presentation and PC differentiation in GCB-DLBCL, regardless of the *EZH2* mutation status. We observed similar transcriptional signatures after treatment with HDAC3i alone, with the addition of 5-Aza promoting an enhanced differentiation transcriptomic profile, reflected by the upregulation of the UPR.

In summary, we demonstrate the efficacy of a new epigenetic therapy for DLBCL consisting of 5-Aza plus HDAC3i. Our data aligns with recent pre-clinical and clinical studies describing promising results with new combinatorial therapies based on HMA. In the case of R-CHOP, 5-Aza priming reprograms chemoresistant DLBCL cells to become sensitive to the immunochemotherapy (8,9). In addition, it has been shown that Decitabine improves the activity of the BCL2 inhibitor Venetoclax, increasing mitochondrial apoptotic priming and BCL-2 dependency of DLBCL cells (35). These results suggest that HMA have the ability to potentiate the anti-lymphoma effects of the other therapeutic agent of the combination. In our study, we observe an increase in the number of H3K27ac peaks at promoters after 5-Aza treatment, which was added to the increase in acetylation mediated by HDAC3 inhibition and resulted in the stronger transcriptional activation of differentiation genes. This effect could be due to the rearrangement of repressive histone marks induced by 5-Aza. In this regard, it has been shown that high doses of 5-Aza reorganizes the patterns of H3K27me3 or H3K9me3 in human embryonic kidney 293 (HEK293) cells (36). Further investigation of the epigenetic changes induced by the combinatorial therapy as well as additional studies on the efficacy of this treatment against primary lymphoma cells will help to determine the translational potential of combining 5-Aza and HDAC3 inhibition to treat DLBCL.

## METHODS

### DLBCL cell lines

The human DLBCL cell line MD901 were grown in medium containing RPMI-1640, 10% FBS, 1% HEPES, 1% glutamine, and 1% penicillin/ streptomycin. MD901 was provided by Jose Angel Martinez-Climent (CIMA, Pamplona, Spain). The DLBCL cell line OCI-Ly7 was grown in Iscove’s medium supplemented with 10% FBS and 1% penicillin/streptomycin. OCI-Ly7 was purchased from the German Collection of Microorganisms and Cell Cultures GmbH (DSMZ). Cell line authentication testing was performed by DNA genotyping, performed by the Australian Genome Research Facility (AGRF; Melbourne, Australia) using the Promega GenePrint 10 System. Short tandem repeat testing was performed, and the genetic profile obtained was compared with the established cell line profile. All cell lines were cultured at 37°C in a humidified atmosphere of 5% CO2. Cell lines were also routinely tested for *Mycoplasma* contamination.

### Culture of PBMC

Human peripheral volunteers were obtained from healthy donors with informed consent following the Declaration of Helsinki according to recommendations of an independent scientific review board. The project has been validated by The Etablissement Français du Sang, the French national agency for blood collection (13-PP-11). Blood samples were collected using ethylene diamine tetraacetic acid (EDTA)–containing tubes. Mononucleated cells were first isolated using Ficoll Hypaque (Eurobio, CMSMSL0101) and cultured in RPMI 1640 medium with glutamax-I (Life Technologies) supplemented with 10% FBS.

### *In vitro* drug treatment

The selective HDAC3 inhibitor (KDAc-001) was kindly provided by KDAc Therapeutics; Panobinostat (LBH589, S1030) and Vorinostat (SAHA, S1047) were obtained from Selleck Chem; 5-Azacytidine (5-Aza) was purchased from Sigma (A2385). For *in vitro* experiments, small molecules were reconstituted to a final stock concentration of 10mM in dimethylsulfoxide (DMSO; HDAC3i, Panobinostat and Vorinostat) or phosphate-buffered saline (PBS; 5-Aza) prior to serial dilution. After seeding (day 0), DLBCL or PBMC cells were treated with 5-Aza daily from day 0 to day 4, and with HDACi or DMSO on days 2 and 4. To calculate synergy, cells were exposed to a dose curve of each drug alone or their combination in a constant ratio. Then, we used CompuSyn software (Biosoft) to plot dose-effect curves.

### Mouse xenografts

The Peter MacCallum Cancer Centre Animal Ethics Committee approved all in vivo procedures. All experimental mice were maintained under specific pathogen-free conditions. For transplantation of lymphoma cells, cohorts of 6-to 8-week-old female NSG mice (bred in house PMCC) were inoculated via intraperitoneal injection with 0,1x10^6^ low-passage OCI-Ly7-Luc2-Cherry cells. Whole-body bioluminescent imaging (BLI) was assessed with an IVIS100 imaging system (Caliper LifeSciences). Mice were injected intraperitoneally with 50 mg/kg D-luciferin (Caliper LifeSciences), anesthetized with isoflurane, and imaged for 2 minutes after a 15-minute incubation following injection. Treatment was initiated once lymphoma cells were clearly visible by bioluminescence, 3-4 weeks after transplantation. For *in vivo* formulation, HDAC3i was dissolved in 0,5% methylcellulose; 0.2% Tween-80 at 2mg/ml and 5-Aza was dissolved in PBS at 0,1mg/ml. Mice were treated daily with 1mg/kg 5-Aza (or vehicle) by intraperitoneal (i.p.) injection for 1 week followed by 3 weeks of treatment with 1mg/kg 5-Aza (or vehicle) daily in combination with 25mg/kg HDAC3i (or vehicle) daily via oral gavage. GraphPad Prism 8 was used for Kaplan–Meier visualization and statistical calculations (Mantle-Cox log-rank test).

### Flow cytometry

Cell viability was determined by propidium iodide (PI) staining. For the analysis of apoptosis, DLBCL cells were assessed for Annexin-V (Biolegend, 422201) and PI double positivity. For CellTrace Violet (CTV) assays, tumor cells were resuspended in PBS (1e7 cells/mL) and labeled with 5mM CTV reagent (Thermo-Fisher Scientific, C34557) in a 37C water bath for 10min. Unbound dye was subsequently quenched through the addition of 5 volumes of ice-cold cell culture media. The cells corresponding to the main peak of Far Red positivity were sorted, and cultured in the presence of vehicle or epigenetic drugs. Data was collected on a FACSCanto II, LSR II or LSR Fortessa flow cytometer (BD Biosciences) and analyzed using FlowJo software (Tree Star).

### Western blot

Protein lysates were prepared using cell lysis buffer (150mM NaCl, 50mM HEPES, 1,5mM MgCl2, 1mM EDTA, 10% glycerol) supplemented with Triton and the cOmplete™ protease inhibitor cocktail (EDTA-free; Roche). Lysates were subjected to SDS-PAGE and transferred to PVDF membrane (Millipore). Membranes were blocked in skim milk or BSA and probed overnight with the following antibodies: Anti-Caspase 3 (Cell Signaling Technology, Cat# 9662); anti-HSP90 (Santa Cruz, Cat# sc-13119). Filters were then washed, probed with either anti-mouse HRP or anti-rabbit HRP (Cell Signaling Technology) secondary antibodies. Signals were visualized using the LAS-4000 digital Imaging System (Fujifilm Life Science).

### Quantitative RT-PCR

Total RNA from DLBCL cells was isolated using TRIzol Reagent (Invitrogen) followed by purification using the Direct-zol RNA Miniprep kit (Zymo Research) following manufacturer’s protocol. Synthesis of cDNA was performed following the standard protocol from the High-Capacity cDNA Reverse Transcription Kit (Applied Biosystems). Quantitative RT-PCR was performed using SYBR Green (Applied Biosystems) method in a 96-well format StepOne Plus (Applied Biosystems). For quantification, the C_T_ values were obtained and normalized to the C_T_ values of *Gapdh* gene. Fold changes in expression were calculated by the 2^-ββCT^ method. Primer sets used for gene expression analysis can be found in Supplementary Table S5.

### RNA-seq and analysis

DLBCL cells were collected on day 5 after starting the treatment. Total RNA was prepared using TRIzol (Invitrogen). Samples were further purified using the Direct-zol RNA Miniprep kit (Zymo Research). mRNA libraries were generated using the QuantSeq 3’-mRNA Seq Library Prep Kit for Ilumina (Lexogen) and sequenced in a NextSeq500 sequencer SE75bp (Illumina). Sequencing files were demultiplexed (Bcl2fastq, v2.17.1.14) and QC was performed on FASTQ files using FASTQC (v0.11.6). Sequencing reads were trimmed (cutadapt v2.1) and aligned to the hg38 human reference genome using HISAT2 (v2.1.0). Read counting across genomic features was performed using FeatureCounts (Subread, v2.0.0) and differential gene expression analysis was performed using Voom-Limma in R (v3.46.0). Gene set enrichment analyses were performed using the Broad Institute GSEA software (37).

### ChIP-seq

25 million DLBCL cells collected on day 5 after starting the treatment were fixed with 0,5% formaldehyde for 10min and quenched with addition of 1.25 mol/L glycine. Cells were lysed in nuclear extraction buffer 3 times. Nuclei were resuspended in sonication buffer and sonicated with a S220 Focused-ultrasonicator (Covaris) (duty cycle 10%; intensity 4; cycle/burst, 200; 6 cycles). Samples were cleared by centrifugation at 12,000*g* for 20 minutes, and 1 volume of dilution buffer was added to cleared chromatin; 1% chromatin was taken as input. Immunoprecipitation was performed overnight at 4°C with rotation with anti-H3K27Ac (Abcam ab4729). Immunoprecipitated samples were captured by incubation with 50μL Dynabeads Protein G (Life Technologies) blocked with 0.1% BSA for 3 hours. Beads were then washed twice each with wash buffer 1, wash buffer 2, wash buffer 3, and TE buffer. DNA was eluted with 150μL elution buffer for 1hr and reverse cross-linked overnight. DNA product was purified using Zymo ChIP DNA Clean and Concentrator Kit. Libraries were generated using the NEBNext Ultra II DNA library prep kit from NEB. For sequencing, libraries were pooled and sequenced on a NovaSeq X Plus Series PE150bp (Illumina), and 15 to 20 million reads were generated per sample. The downstream bioinformatic analysis was performed by Marbyt.

### RRBS

Genomic DNA from DLBCL cells was isolated using the Monarch gDNA purification kit (New England Biolabs). DNA methylation library preparation was performed by Novogene, including sequencing on the NovaSeq X Plus Series PE150bp (Illumina), and 35-45 million reads were obtained per sample. The downstream bioinformatic analysis was performed by Marbyt.

### MAC-seq

At day 4 of treatment, 20000 cells from each well were aliquoted into a separate 96-well plate, washed in ice-cold PBS twice and centrifuged (1400 rpm at 4 °C for 4 min). Supernatant was removed, and cell pellets were frozen at −80 °C. Library preparation was performed by the addition of 15 µl lysis buffer into each well of a 96-well plate containing cell pellets and incubated at room temperature for 15 min under agitation (900 rpm). 12.5 µl of cell lysate was transferred into each well of a new 96-well plate previously prepared with 1 µl of 10 nM well-specific RT MAC-seq primer and 7.5 µl RT mix; the RT mix contains a TSO primer and external ERCC RNAs for normalization. The mixture was incubated for 2 hr at 42 °C to create well-barcoded full-length cDNA, and then all the wells of a plate were combined into a single tube. Concentration and clean-up were done with DNA Clean & ConcentratorTM-100 (Zymo Research), and RNAClean XP (Beckman Coulter) and each plate were eluted in 22 µl nuclease-free water. The purified cDNA was pre-amplified with KAPA HiFi HotStart ReadyMix (Roche), and MAC-seq PreAmp PCR primer and the quality checked on a D5000 Screentape (TapeStation, Agilent). One barcoded library was prepared per plate using TD buffer and TDE1 enzyme (Illumina) for tagmentation and KAPA HiFi HotStart Ready Mix (Roche) and custom primers (MAC-seq P5 PCR and MAC-seq Indexing Mix) for amplification. Libraries were purified with DNA AMPure XP (Beckman Coulter), quality checked on a DNA1000 tape (TapeStation, Agilent) and quantity verified by qPCR. Two indexed libraries were sequenced on a NextSeq 500 instrument (Illumina) using a custom sequencing primer (MAC-seq Read primer) and a High Output Kit v2.5 75 Cycles (Illumina) with paired-end configuration (25 base pairs for read 1 and 50 base pairs for read 2). Paired-end reads were demultiplexed using bcl2fastq (v2.17.1.14) and resulting FASTQ files were quality checked using fastqc (v0.11.6) and read 2 (R2) was trimmed 15 bp from the 5′ end to remove primer bias using cutadapt (v2.1; −u 15). R2 FASTQ files of paired-end reads were demultiplexed according to well barcodes (Supplementary Table S6) and filtered for PCR duplicates using Unique Molecular Identifiers (UMIs), both present in read 1 (R1) using the scruff 9 R (v4.0.2) package dumultiplex function (bcStart = 1, bcStop = 10, bcEdit = 0, umiStart = 11, umiStop = 20, keep = 35, minQual = 20, yieldReads = 1e + 06). R2 FASTQ files were then mapped to the GRCh38/hg38 genome and ERCC sequences using alignRsubread (unique = FALSE, nBestLocations = 1, format = “BAM”) and resulting BAM files were used to count unique R2 reads mapping to exonic genomic intervals and ERCC sequences using a combined hg38/ERCC GTF file with countUMI (umiEdit = 0, format = “BAM”, cellPerWell = 1. Both functions are from the scruff R package. Gene expression counts were normalized to library size. Subsequent count processing was performed using the Seurat R package (v3.2.1) 10, where lowly expressed genes were filtered, and counts were normalized for latent variables including plate, well row and column, using the SCTransform function. SCTransformed scaled gene RNA expression values were then used for PCA, where shared-nearest-neighbours (SNN) network was calculated using the top 10 Principal Components with the FindNeighbours function using default parameters. Drug-treatment clusters were subsequently identified with the Louvain algorithm using a resolution parameter of 2. Uniform Manifold Approximation and Projection (UMAP) values were also calculated using the top 10 Principal Components with the RunUMAP function using default parameters. Differential gene testing relative to treatment controls was performed using a hurdle model (MAST) and a logFC threshold of 0 with the FindMarkers Function. Area Under the Curve (AUC) scores for each drug treatment and gene lists indicated was calculated using all expressed genes with the R AUCell Package (v0.10.0). *ggplot2* (version 3.2.1) was used to visualize the data.

### Statistics

Statistical differences between the means of two data groups was determined by using two-tailed unpaired Student’s *t* test, and *p* < 0.05 were considered significant. Multiple group comparisons were performed using ANOVA, *p* < 0.05 were considered significant.

## Data availability

All sequencing datasets are available on the Gene Expression Omnibus (GEO) under accession number.

## ACKNOWLEDGMENTS

We thank staff from the Animal Facility and Flow Cytometry Facility of the C3M, the Animal Facility, Molecular Genomics Core, Flow Cytometry Facility, Victorian Center for Functional Genomics (VCFG) and Bioinformatics Consulting Core (BCC) of the Peter MacCallum Cancer Centre, members of the Johnstone and Melnick laboratories for useful discussions, and Ana Steiner, Mark Inston and Alison Jones for their important contribution as consumer advocates. This research has been funded by the Leukemia & Lymphoma Society-Snowdome Foundation-Leukaemia Foundation Translational Research Program (TRP, grant #6605-20), the Australian National Health and Medical Research Council (NHRMC, grant #2021_GNT2011217), the Fondation ARC pour la Recherche sur le Cancer (Recruiting international leaders in Oncology 2023), La Ligue contre le cancer (Allocations de recherche Doctorals 2024, C. Calaud) and the French Government (National Research Agency, ANR) through the “Investments for the Future” IDEX UCAJedi **ANR-15-IDEX-01**.

## Notes

### Competing Interest Statement

The authors have declared no competing interest.

