## Supplementary data for "Enhanced efficacy of a specific HDAC3 inhibitor in combination with 5-Azacitidine against diffuse large B-cell lymphoma"

[6]Equipe Leader Fondation ARC 2023.

* These authors contribute equally.

### Senior author.

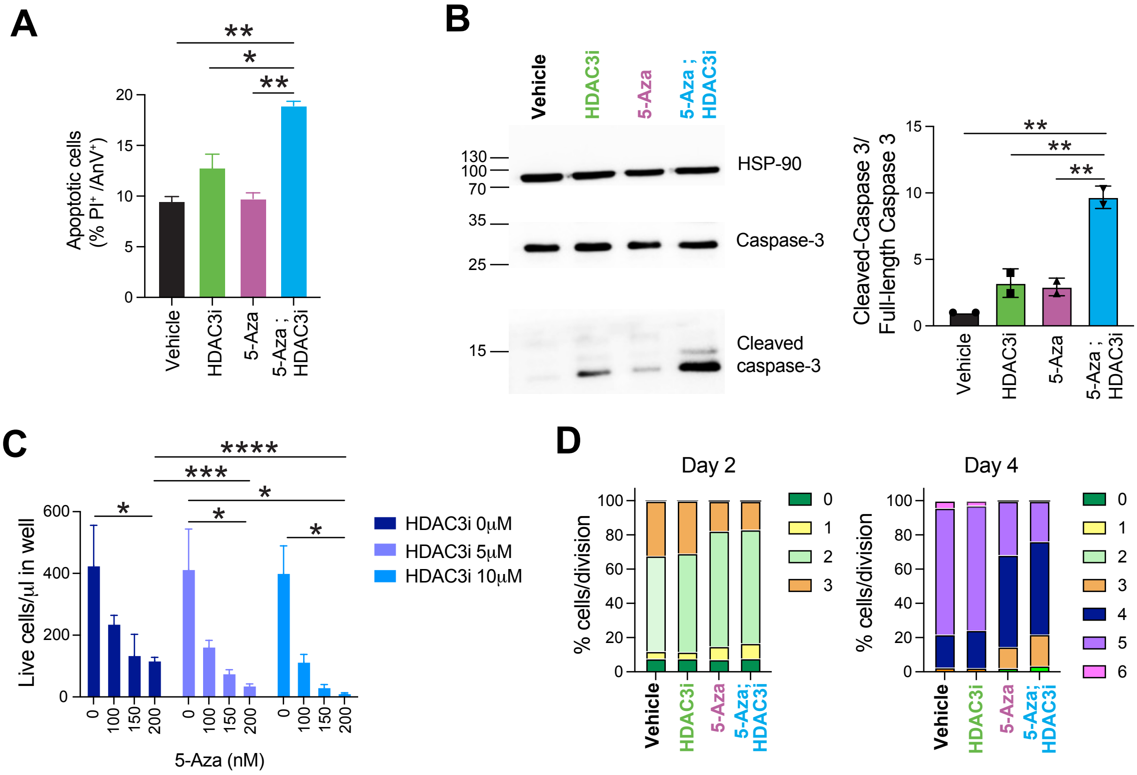


**Supplementary Fig. S1: 5-Aza and HDAC3i synergize to increase cell death and reduce proliferation in DLBCL cells.** (**A**) Representative density plot and quantification of Annexin $V^+$ $PI^+$ cells at day 7 after treatment of OCI-Ly7 cells with 5-Aza and HDAC3i. (**B**) Representative western blot of full-length Caspase 3 and cleaved-Caspase 3, and quantification relative to control at day 5 after treatment of OCI-Ly7 cells with 5-Aza and HDAC3i. (**C**) Total cell number of OCI-Ly7 cells at day 7 after treatment with 5-Aza and HDAC3i. (**D**) CTV assay at day 2 and day 4 after treatment of OCI-Ly7 cells with 5-Aza and HDAC3i.

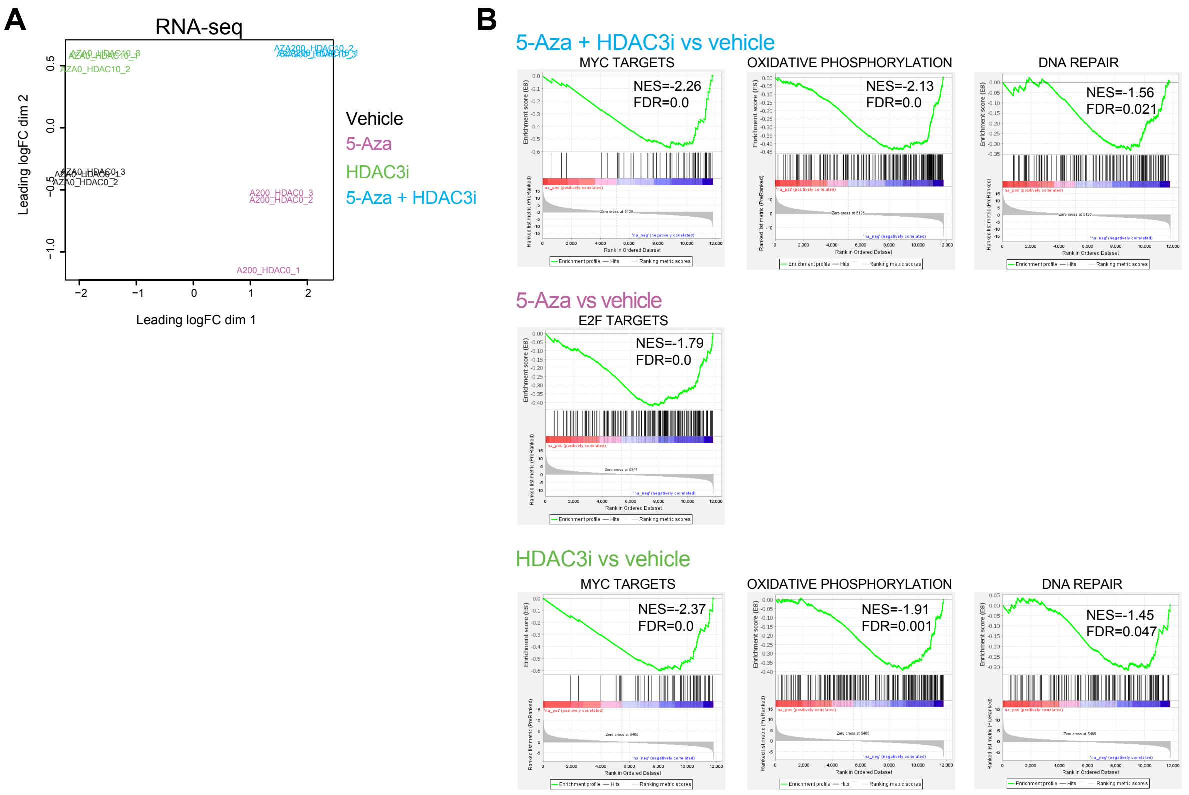


**Supplementary Fig. S2: RNA-seq analysis of DLBCL treated with 5-Aza and HDAC3i.** (**A**) Principal component analysis of RNA-seq data. (**B**) Gene set enrichment analysis using the Hallmark gene sets showing negatively enriched signatures.

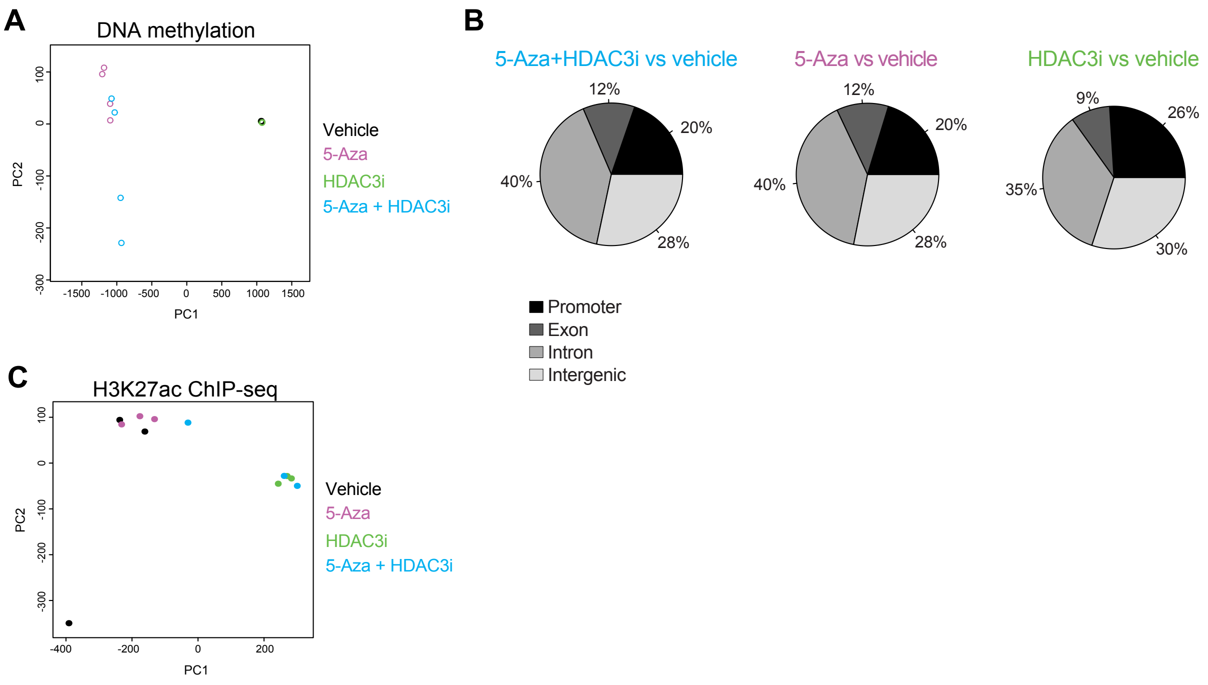


**Supplementary Fig. S3: Multi-omics analysis of DLBCL treated with 5-Aza and HDAC3i.** (**A**) Principal component analysis of RRBS data. (**B**) Genomic distribution of the DMC between 5-Aza+HDAC3i, 5-Aza or HDAC3i and vehicle. (**C**) Principal component analysis of H3K27ac ChIP-seq data.

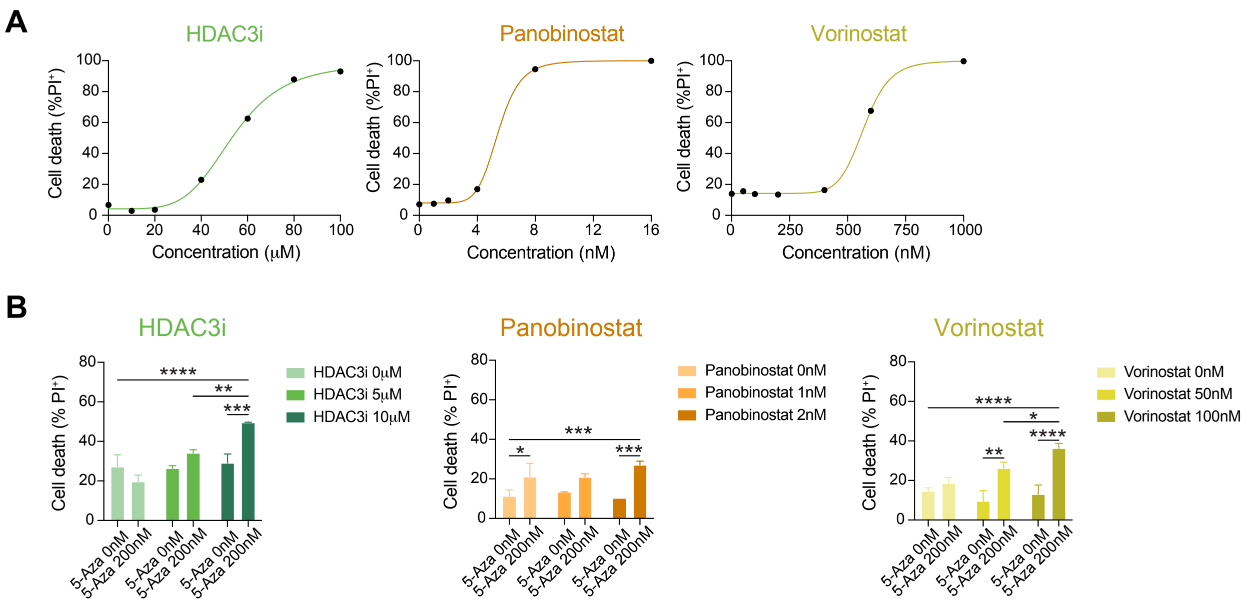


**Supplementary Fig. S4: Comparison of 5-Aza+HDAC3i vs 5-Aza+pan-HDACi.** (**A**) Dose curves of HDAC3i, Panobinostat and Vorinostat on MD901 cells analyzed at day 7. (**B**) Percentage of $PI^+$ cells at day 7 after treatment of MD901 cells with vehicle or 5-Aza in combination with HDAC3i, Panobinostat or Vorinostat in a 96-well plate format.

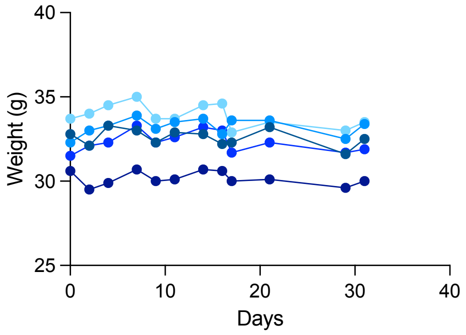


**Supplementary Fig. S5:** Weights of the lymphoma-bearing mice cells throughout the duration of the treatment with 5-Aza+HDAC3i.
